# Hyperactive chemotaxis contributes to anti-TNFα treatment resistance in inflammatory bowel disease

**DOI:** 10.1101/2021.08.15.456400

**Authors:** Tung On Yau, Jayakumar Vadakekolathu, Gemma Ann Foulds, Guodong Du, Christos Polytarchou, Benjamin Dickins, Sergio Rutella

## Abstract

**Background & Aims:** Anti-tumour necrosis factor-alpha (anti-TNFα) agents have been used for inflammatory bowel disease (IBD), however, it has up to 30% non-response rate. Identifying molecular pathways and finding reliable diagnostic biomarkers for patient response to anti-TNFα treatment are clearly needed.

**Methods:** Publicly available transcriptomic data from IBD patients receiving anti-TNFα therapy was systemically collected and integrated. *In silico* flow cytometry approaches and MetaScape were applied to evaluate immune cell populations and to perform gene enrichment analysis, respectively. Genes identified within enrichment pathways validated in neutrophils were tracked in an anti-TNFα-treated animal model (with lipopolysaccharide (LPS)-induced inflammation). The receiver operating characteristic (ROC) curve was applied to all genes to identify the best prediction biomarkers.

**Results:** A total of 449 samples were retrieved from control, baseline and after primary anti-TNFα therapy or placebo. No statistically significant differences were observed between anti-TNFα treatment responders and non-responders at baseline in immune microenvironment scores. Neutrophils, endothelial and B cell populations were higher in baseline non-responders and chemotaxis pathways may contribute to the treatment resistance. Genes related to chemotaxis pathways were significantly up-regulated in LPS-induced neutrophils but no statistically significant changes were observed in neutrophils treated with anti-TNFα. Interleukin 13 receptor subunit alpha 2 (*IL13RA2*) is the best predictor (ROC: 80.7%, 95% CI: 73.8% - 87.5%) with a sensitivity of 68.13% and specificity of 84.93%, and significantly higher in non-responders compared to responders (*p* < 0.0001).

**Conclusions:** Hyperactive chemotaxis influences responses to anti-TNFα treatment and *IL13RA2* is a potential biomarker to predict anti-TNFα treatment response.

## Introduction

Inflammatory bowel disease (IBD) is an idiopathic and relapsing-remitting chronic inflammatory disorder characterised by a susceptible genetic background, causing immunological dysfunction and intestinal microbiome dysbiosis.^1^ The long-standing mucosa inflammation destruct tight junctions, induces intestinal barrier injury and permeability and increase the incidence of colonic neoplasia.1 It is estimated that the prevalence of IBD exceeds 0.3% in North America, Oceania, and many countries in Europe.^2^ With the incidence rising in the newly industrialised countries, including Brazil and Taiwan,^3^ thus, IBD places a large burden on public health services and healthcare economies.

Tumour necrosis factor-alpha (TNFα) is a pleiotropic cytokine that participates in several pathological processes in IBD and is recognised as a pro-inflammatory cytokine. The production of biologically active homotrimer TNFα originally from the precursor TNFα through TNF-converting enzyme (TACE) proteolysis from the soluble TNFα.^4^ TNFα activity is mediated through binding to the TNF receptors I and II (TNFRI and TNFRII).^5^ This binding activates immune cells response and pro-inflammatory cytokine and chemokine productions, such as IL-1, IL-6, IL-8 and RANTES. It also increases the expression of adhesion molecules, production of matrix metalloproteinase and induction of apoptosis.^6^ The use of anti-TNFα compounds such as full monoclonal IgG1 antibodies (infliximab and adalimumab), pegylated anti-TNFα F[ab’]2 fragment (certolizumab) and IgG1қ monoclonal antibody derived from immunising genetically engineered mice with human TNFα (golimumab) have been approved for IBD patients^7^, including Crohn’s disease (CD) and ulcerative colitis (UC) and IBD unclassified (IBD-U)^8^.

Although CD and UC are the distinct subtypes of IBD, these diseases present a certain level of similarities, including symptoms, pathological features, immune response, risk factors and the biological pathways producing TNFα.^9^ In addition, studies found that up to 3% of CD patients will be reclassified as UC and *vice versa* after their primary diagnosis, 5 – 15% of IBD patients classified as IBD-U and a small portion of UC patients is later changed to CD or IBD-U.^10^ More importantly, up to 30% of patients do not respond to anti‐TNFα blockers^1,11^ and the use of vedolizumab (Anti-IL-12/23) and ustekinumab (anti-integrin), may be efficacious in many patients that failed anti-TNFα therapy.^12^ Thus, there is a clear need to identify potential anti-TNFα treatment pathways in overall IBD patients with a view to better targeting anti-TNFα treatment to more responsive cohorts and to minimise the adverse anti-TNFα treatment effects.

## Methods

### Search strategy, data collection and integration

A searching strategy for publicly available datasets related to IBD patients received anti-TNFα therapy was designed for the NCBI Gene Expression Omnibus (GEO) database dated December 31, 2020, using the keywords, “TNF”, “Tumor Necrosis Factor”, “anti-TNF”, “anti-Tumor Necrosis Factor”, “Infliximab”, “Adalimumab”, “Golimumab”, “inflammatory bowel disease”, “IBD”, “ulcerative colitis”, “UC”, “Crohn Disease” and “CD”. The included datasets have to meet the following inclusion criteria: (1) colonic sample from IBD patients, (2) transcriptomic data, (3) raw data are available, (4) anti-TNFα treatment response status, (5) publicly accessible (6) each of the original studies obtained approval from their local ethics committee and had written, informed patient consent. Sample exclusion criteria were as follows: (1) subjects receive therapy other than anti-TNFα, (2) overlapped subjects, (3) colonic samples other than large intestine, (4) transcriptomic data other than Affymetrix, (5) the post-treatment time point being over three months.

The eligible raw microarray datasets were collected and subjected to background correction, normalisation, and summarisation using the Robust Multichip Average (RMA) algorithm using Affy package version 1.66.0 individually.^13^ Mean value of multiple probe sets representing the same gene was calculated. Next, the ComBat function from sva package version 3.36.0 was implemented on the datasets to eliminate the study-specific batch effects.^14,15^

### Composition of immune cells and immune-related scores evaluation

The evaluation of immune microenvironment scores and immune-stroma cells population are calculated using xCELL^16^ and ESTIMATE (Estimation of Stromal and Immune cells in MAlignant Tumor tissues using Expression data)^17^ algorithms. The immune cell types were evaluated from gene expression profile using 5 different algorithms, including CIBERSORT,^18^ EPIC,^19^ MCP-Counter,^20^ xCELL and Deconvolution-To-Estimate-Immune-Cells (DTEIC)^21^. Each of the algorithms was developed using their in-house or publicly immune cells expression data and different statistical learning approaches. For instance, DTEIC utilised ε-support vector regression (ε-SVR) and CIBERSORT applied linear support vector regression (SVR)^18,21^; MCP-Counter is a single sample scoring system while xCELL requires heterogenous dataset^16,20^; ESTIMATE utilises single-sample Gene Set Enrichment Analysis (ssGSEA) to rank samples on the expression of two different 141-gene sets, and xCELL is based on the sets of cells values calculated from its algorithm^16,17^.

The scores/cells population from EPIC version 1.1, MCP-Counter version 1.2.0 and ESTIMATE version 1.0.13 were performed under R version 4.0.0 programming environment and DTEIC was operated under Python 3.7 programming environment. Both CIBERSORT and xCELL were calculated using their corresponding online tools. The source code can be found on the corresponding authors’ GitHub page.

### Functional Enrichment Analysis

Identification of differentially expressed genes (DEGs) between responders and non-responders were calculated using limma package version 3.22.3,^22^ the threshold for the DEGs has a Benjamini-Hochberg adjusted *p*-value < 0.05 with absolute log 2-fold change ≥ 0.75. EnhancedVolcano package version 1.6.0 was applied for Volcano plot.^23^ Heatmap was generated by using pheatmap package version 1.0.12.^24^ All the packages are applied within the R programming environment. The differentially over-expressed genes were utilised for the pathway enrichment analysis using Metascape (http://metascape.org),^25^ a gene enrichment tool for understanding from previously pre-defined gene sets in different enriched biological themes, including GO terms, Kyoto Encyclopedia of Genes and Genomes (KEGG), Reactome, BioCarta and MSigDB. For each gene inputted into the server, the enrichment score was calculated and clustered to match biological signalling pathways. Visualisation of the selected pathways utilised Cytoscape version 3.8.0.

### Lipopolysaccharide-induced inflammation in neutrophils

To further confirm the outcomes from the functional enrichment analysis, experimental neutrophils data from *Macaca mulatta* was applied. Briefly, neutrophils were collected from the target site (at approximately 130 days of gestation). Inflammation was subsequently induced at this site via lipopolysaccharide (LPS) treatment. Subjects were either treated or not treated with adalimumab at 3 hours and 1 hour before LPS, with samples taken at 16 hours post LPS.^26^ The original study obtained approval from their local ethics committees. The raw count data retrieved from GSE145918 and normalised the values were calculated using per million reads mapped (CPM) from the count matrix used edgeR version 3.32.0 and were log_2_ + 1 transformed under R programming environment.

### Statistics

Statistical analysis was performed using R version 4.0.0. The pROC package version1.16.2 in R programming environment was applied to conduct receiver operating characteristic (ROC) curve analysis to evaluate diagnostic accuracy. The statistical significance was evaluated using a Mann-Whitney test, Benjamini and Hochberg adjustment was applied for the IBD treatment data. Differences were considered statistically significant at a *p*-value of < 0.05, and < 0.05, < 0.01, < 0.001 and < 0.0001 are indicated with one, two, three and four asterisks, respectively.

## Results

### Characteristics of studies included in the analysis

After the keyword searching, removal of ineligible and overlapped datasets from the total of 182 records, 5 transcriptomic data, including GSE16879: from the University Hospital of Gasthuisberg, Belgium with ClinicalTrials.gov number NCT00639821;^27^ GSE23597: the multicentre, randomised, double-blind, placebo-controlled ACT-1 study between March 2002 and March 2005 with ClinicalTrials.gov number NCT00036439;^28^ GSE52746: the colonic samples collected between November 2010 and November 2013 from the Department of Gastroenterology, Hospital Clinic of Barcelona, Spain;^29^ GSE73661, UC samples collected from two phase III clinical trials of Vedolizumab (VDZ) - GEMINI I and GEMINI LTS at Leuven University Hospitals, Belgium, the dataset included patients received anti-TNFα blockers;^30^ GSE92415: the PURSUIT golimumab study conducted in multi-centres, with ClinicalTrials.gov number NCT01988961.^31^ The five eligible microarray datasets were normalised, combined and batch effects corrected (**Supplementary Figure 1**). Eventually, a total of 374 samples, with 17,771 common gene symbols were included in this study (**Table 1**).

**Table 1.** Summary of the included transcriptomic studies from large intestinal tissues in IBD patients.

| GSE No. | Affymetrix Platform | Anti-TNFα drug | Study Location | IBD | Pre-Treatment | | Post-Treatment | | Control | Time Point (week) |
| --- | --- | --- | --- | --- | --- | --- | --- | --- | --- | --- |
|  |  |  |  |  | R | NR | R | NR |  |  |
| 16879<sup>32</sup> | Human Genome U133 Plus 2.0 | infliximab | Belgium | UC | 8 | 16 | 8 | 16 | 6 | 4–6 |
|  |  |  |  | CD | 12 | 7 | 11 | 7 |  |  |
| 23597<sup>33</sup> | Human Genome U133 Plus 2.0 | infliximab | USA | UC | 25 | 7 | 20 | 7 | - | 8 |
| 52746<sup>34</sup> | Human Genome U133 Plus 2.0 | Infliximab or adalimumab | Spain | CD | 6 | 1 | 7 | 5 | 17 | 12 |
| 73661<sup>35</sup> | Human Gene 1.0 ST | infliximab | Belgium | UC | 8 | 15 | 8 | 15 | 12 | 6/12 |
| 92415<sup>36</sup> | HT HG-U133+ PM | Golimumab | USA | UC | 32 | 27 | 29 | 21 | 21 | 8 |
| **Over-all = 374** |  |  |  | UC | 73 | 65 | 65 | 59 | 56 |  |
|  |  |  |  | CD | 18 | 8 | 18 | 12 |  |  |
|  |  |  |  | **Total** | **91** | **73** | **83** | **71** |  |  |

CD: Crohn's Disease; UC: Ulcerative Colitis; R: responder; NR, non-responder. TNFα , tumour necrosis factor alpha.

### Immune microenvironment cells population are significantly higher in non-responders

Firstly, the immune microenvironment scores from both ESTIMATE and xCELL identified the baseline anti-TNFα treatment non-responders are significantly higher in compared to the responders (ESTIMATE: *p* < 0.0001, xCELL: *p* = 0.0003) (**Figure 1A–B**). The TNFα treatment responders showed a significant drop after their treatments (ESTIMATE: *p* < 0.0001, xCELL: *p* = 0.0004) while no significant changes in the non-responders (ESTIMATE: *p* = 0.0650, xCELL: *p* = 0.11) (**Figure 1A–B**). The immune and stoma scores (the two calculation factors for the immune microenvironment) are also significantly higher in baseline anti-TNFα treatment non-responders compared to the responders (**Supplementary Figure** 2A-D).

**Figure 1.**
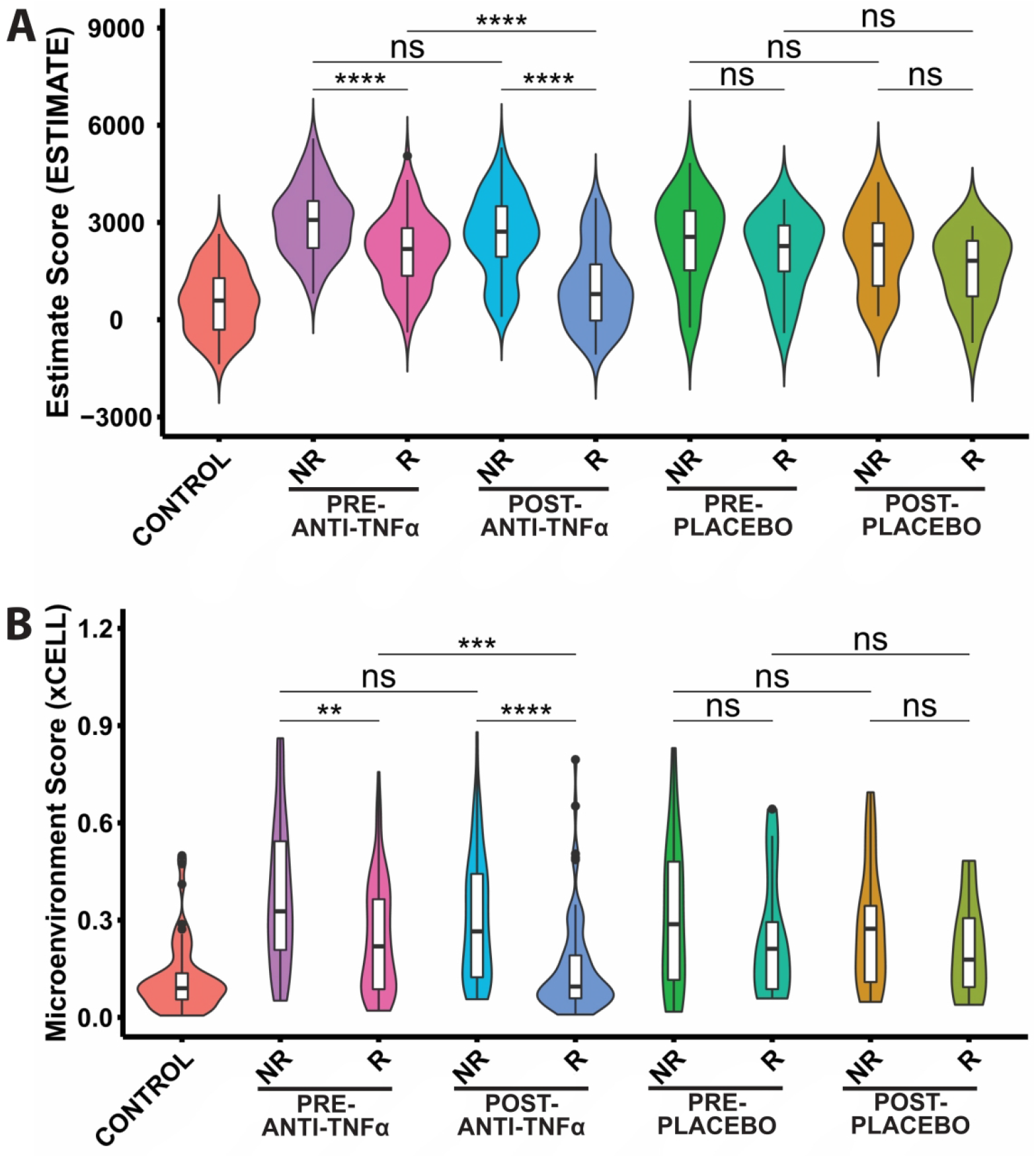
Microenvironment scores are significantly higher on baseline non-responders compared to the responders. Immune microenvironment scores evaluated via (A) ESTIMATE and (B) xCELL algorithms. NR: non-responder; R: responder. The y-axes are the relative immune microenvironment scores from the corresponding algorithms. P-value determines by Mann-Whitney test with Benjamini and Hochberg adjustment. Asterisks denote statistically significant differences (***p < 0.001, ****p < 0.0001).

### Neutrophils, endothelial and B cells are significantly higher in non-responders

To further our understanding of the immune cell-type composition between the treatment responders and non-responders, five different *in silico* flow cytometry approaches, including CIBERSORT, xCELL, EPIC, MCP-Counter and DTEIC were applied (**Supplementary Data 1**). Across the algorithms, neutrophils (MCP-Counter: *p* < 0.0001; xCELL: *p* = 0.0084; CIBERSORT: *p* = 0.0021) (**Figure 2A–C**), endothelial cells (MCP-Counter: *p* = 0.0009; xCELL: *p* = 0.0183; EPIC: *p* = 0.0337) (**Figure 2D–F**) and B cells/B linage (MCP-Counter: *p* = 0.0042; xCELL: *p* = 0.0251; EPIC: *p* = 0.0042) (**Figure 2G–I**) are significantly higher in baseline treatment non-responders compared to the responders. The three-dimensional plots illustrated neutrophils, endothelial cells and B cell from both MCP-Counter and xCELL have positive Pearson correlations with each other using all the eligible data (**Figure 2J–K, Supplementary Figure 3A–B**).

**Figure 2.**
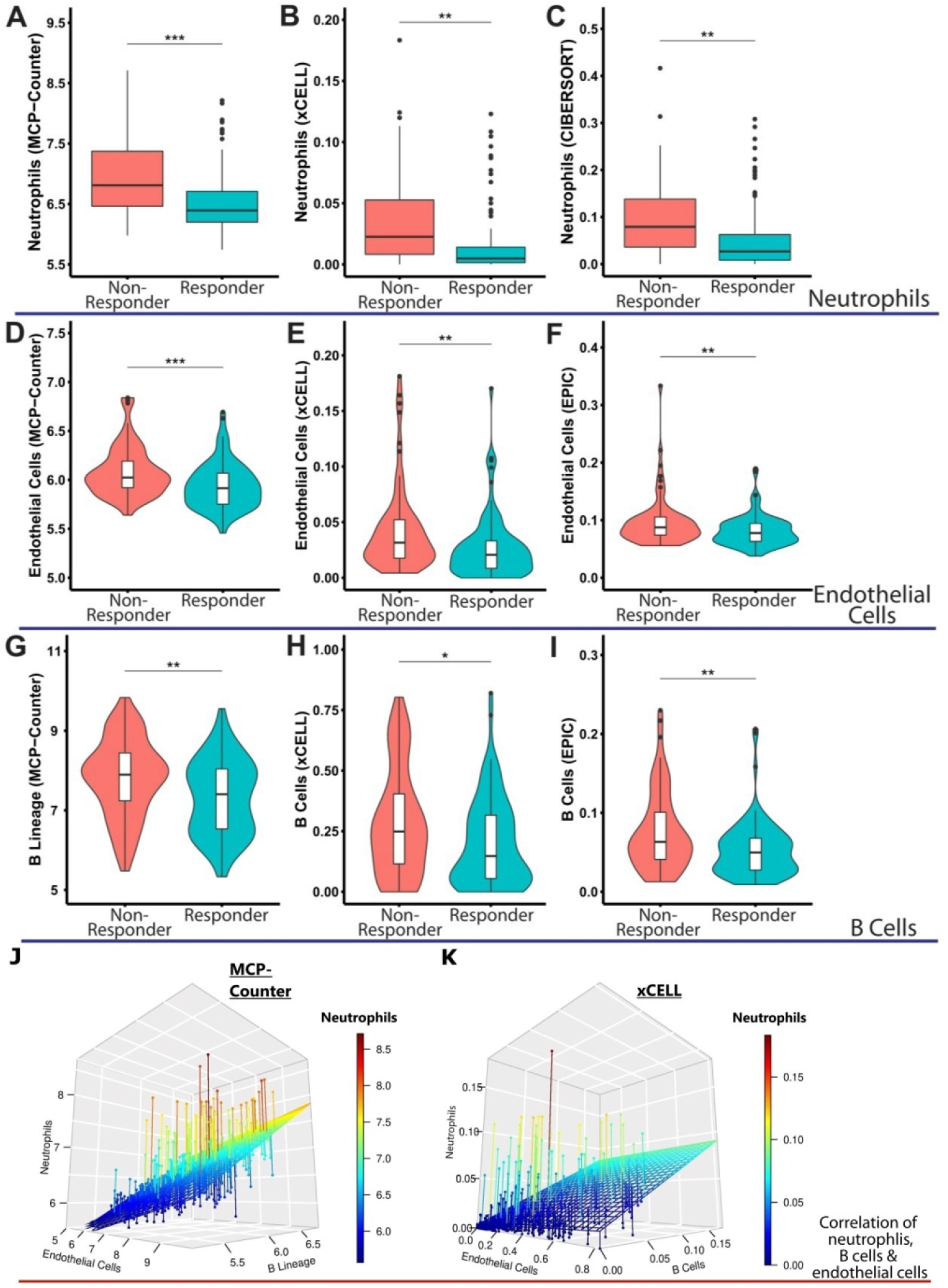
Neutrophils, endothelial cells and B cells are significantly higher on the baseline anti-TNFα treatment non-responders compared to responders. Immune cells population evaluated in five in silico flow cytometry, and (A-C) B cells, (D-F) endothelial cells and (G-I) neutrophils can be recognised in three out of five algorithms. B cells, endothelial cells and neutrophils cells’ populations are higher on baseline anti-TNFα non-responders compared to responders. The three-dimensional plots illustrated (J-K) neutrophils, endothelial cells and B cell populations from MCP-Counter and xCELL algorithms have positive correlations with each other. The y-axes are the relative immune cells population abundance from the corresponding algorithms. NR: non-responder; R: responder. P-value determines by Mann-Whitney test. Asterisks denote statistically significant differences (*p < 0.05, **p < 0.01, ***p < 0.001, ****p < 0.0001).

### Hyperactive chemotaxis contributes to anti-TNFα treatment resistance in inflammatory bowel disease

The pre-treatment anti-TNFα subjects (responder: n = 91 and non-responder: n= 73) were utilised for differentially expressed genes (DEGs) analysis and identified a total of 77 DEGs (upregulated genes = 64, and downregulated genes = 13) (**Figure 3A–B and Supplementary Data 2**). Principal component analysis (PCA) does not have a clear separation between responders and non-responders subjects in the DEGs (**Figure 3C**). The differently up-regulated genes compose of several gene families, including cytokines (*CCL2*, *CCL3*, *CCL4*, *CXCL13*, *CXCL5*, *CXCL6* and *CXCL8*), chemokines (*IL1B*, *IL6*, *IL11* and *IL24*), S100 protein family (*S100A8*, *S100A9* and *S100A12*), selectin (SELE and SELL), matrix metalloproteinases (*MMP1*, *MMP3* and *MMP10*) and formyl peptide receptors (*FPR1*, *FPR2*). Metascape pathways enrichment analysis on the 64 highly expressed genes revealed that GO terms with chemotaxis are commonly found from the outcomes and may have a critical role affecting anti-TNFα treatment (GO:0030595: leukocyte chemotaxis, GO:0002688: regulation of leukocyte chemotaxis, GO:1901623: regulation of lymphocyte chemotaxis and GO:0050918: positive chemotaxis) (**Figure 3D–E, Supplementary Data 3**).

**Figure 3.**
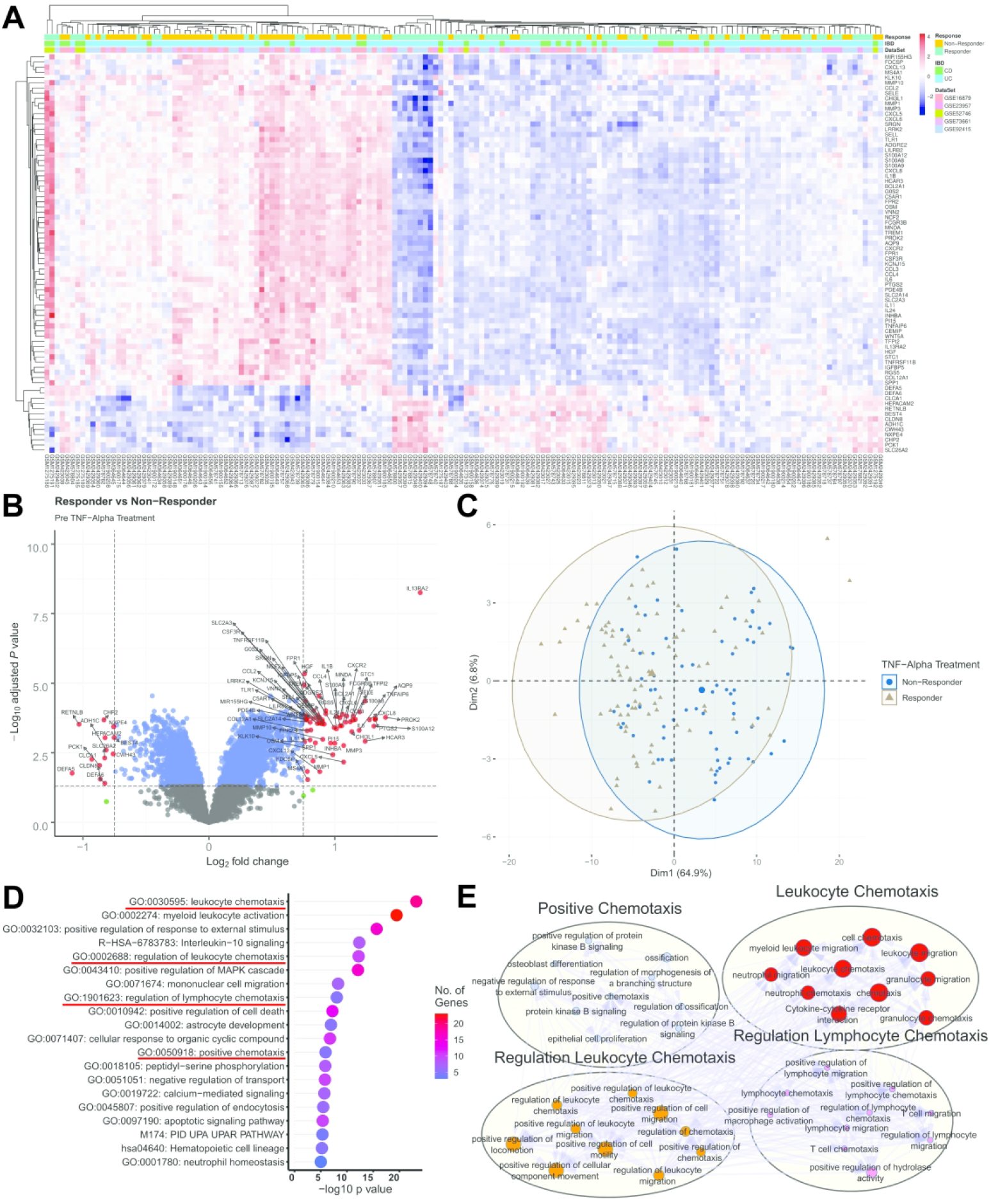
Hyperactive chemotaxis may be involved in anti-TNFα treatment resistance in inflammatory bowel disease. To identify the molecular mechanisms between anti-TNFα blocker responders (n = 91) and non-responders (n= 73), global gene expression analysis was applied from 5 combined and normalised microarray datasets. The differentially expressed genes were identified and presented using (A) heatmap (the relative expression values were z-score transformed), (B) volcano plot and (C) principal component analysis from a total of 64 up-regulated and 13 down-regulated genes based on the adjusted p-value < 0.05 with absolute log 2-fold change ≥ 0.75. (D) The significantly up-regulated genes were utilised for Metascape pathway enrichment analysis from the previously pre-defined gene set. Enriched terms related to the chemotaxis-related pathways are underlined. The y-axis represents the top 20 gene sets category, the x-axis represents -log10 p-value, the colour intensity of the bar represents the number of genes identified in each hallmark category. (E) The four subsets of enriched terms under the chemotaxis-related pathways were selected and visualised using Cytoscape.

### Anti-TNFα blocker does not reduce chemotaxis in LPS-induced inflammation in neutrophils

To demonstrate our finding in chemotaxis, we used the RNA-sequencing data from an animal study.^26^ Briefly, neutrophils isolated from chorio-decidua cells with LPS-induced inflammation were treated with or without adalimumab.^26^ The list of genes from the four chemotaxis enrichment terms matched with the neutrophils data to get the mean expression values of each sample (GO:0030595: leukocyte chemotaxis [31 out of 44 genes], GO:0002688: regulation of leukocyte chemotaxis [18 out of 22 genes], GO:1901623: regulation of lymphocyte chemotaxis [13 out of 16 genes] and GO:0050918: positive chemotaxis [18 out of 21 genes]) (**Supplementary Data 4**). The data process workflow is in **Supplementary Figure 4.** All the enrichment terms related to chemotaxis were significantly higher in the LPS-exposed neutrophils, three out of the four enrichment terms no significant reduction was observed in the anti-TNFα treated group (**Figure 4**).

**Figure 4.**
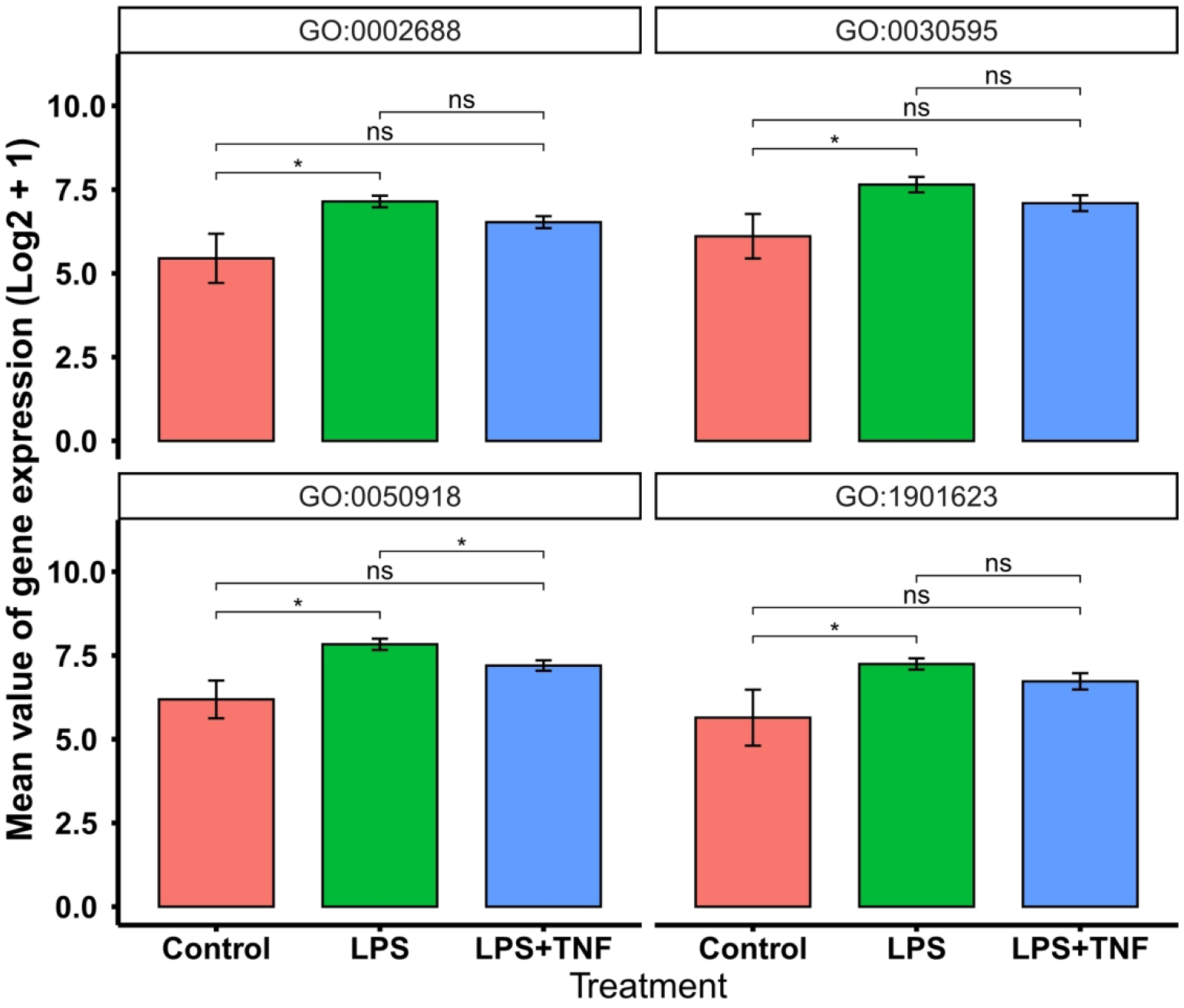
anti-TNFα blocker does not reduce chemotaxis in lipopolysaccharide-induced inflammation in neutrophils. Mean expression level (across samples) of genes matching indicated GO term. LPS, lipopolysaccharide. TNFα, anti-Tumour necrosis factor-alpha. P-value determines by Mann-Whitney t-test. Asterisks denote statistically significant differences (*p < 0.05).

### Interleukin 13 receptor subunit alpha 2 is a diagnostic biomarker predicts TNFα treatment response

In order to find the best potential biomarker to predict anti-TNFα respond IBD patients, ROC curve analysis was applied to all the genes one by one by using a for-loop with pROC package under the R programming environment. Among them, *IL13RA2* has the AUC of 80.7% (95% confidence interval (CI): 73.8% - 87.5%) with the best sensitivity of 68.13% and specificity of 84.93% (**Table 2**, **Figure 5B, Supplementary Data 5**). *IL13RA2* was stand-alone from the volcano plot (log_2_FC:1.678, adjusted *p* < 0.0001) (**Figure 2B**) and the expression of *IL13RA2* is significantly higher in pre-treatment non-responders compared to the pre-treatment responders (*p* < 0.0001). The responders showed a bigger drop after the treatment compared to the non-responders (Responder: *p* < 0.0001, Non-responder: *p* = 0.0037), and the responders restored the expression level to normal control after the treatment (mean ± standard deviation: control: 4.721±1.039, post-treatment, responder: 4.868 ±1.364; *p* = 0.4969) (**Figure 5A**).

**Table 2.** The top 15 genes to predict TNFα treatment response.

| Rank | Gene Name | AUC (95%CI) | Threshold | Best Specificity | Best Sensitivity | LR+ | LR- |
| --- | --- | --- | --- | --- | --- | --- | --- |
| 1 | IL13RA2 | 0.807 (0.738-0.875) | 6.263 | 0.849 | 0.681 | 4.510 | 0.222 |
| 2 | ADGRE2 | 0.786 (0.716-0.857) | 6.079 | 0.795 | 0.703 | 3.429 | 0.292 |
| 3 | ADGRL2 | 0.771 (0.698-0.843) | 6.013 | 0.753 | 0.736 | 2.980 | 0.336 |
| 4 | HGF | 0.768 (0.695-0.841) | 5.598 | 0.753 | 0.725 | 2.935 | 0.341 |
| 5 | TLR1 | 0.764 (0.69-0.838) | 5.95 | 0.836 | 0.637 | 3.884 | 0.257 |
| 6 | NCF2 | 0.757 (0.683-0.831) | 7.696 | 0.671 | 0.769 | 2.337 | 0.428 |
| 7 | RGS5 | 0.757 (0.682-0.832) | 7.105 | 0.767 | 0.681 | 2.923 | 0.342 |
| 8 | FPR1 | 0.756 (0.682-0.83) | 6.947 | 0.836 | 0.626 | 3.817 | 0.262 |
| 9 | TMTC1 | 0.754 (0.68-0.828) | 6.085 | 0.849 | 0.593 | 3.927 | 0.255 |
| 10 | CCL4 | 0.754 (0.679-0.829) | 7.955 | 0.781 | 0.659 | 2.304 | 0.434 |
| 11 | PDE4B | 0.754 (0.679-0.829) | 7.681 | 0.671 | 0.758 | 3.009 | 0.332 |
| 12 | IGFBP5 | 0.754 (0.678-0.829) | 7.185 | 0.712 | 0.725 | 2.517 | 0.397 |
| 13 | SRGN | 0.753 (0.678-0.828) | 9.849 | 0.808 | 0.615 | 3.203 | 0.312 |
| 14 | PAPPA | 0.751 (0.675-0.827) | 5.946 | 0.603 | 0.835 | 2.103 | 0.475 |
| 15 | TMEM71 | 0.749 (0.674-0.824) | 5.672 | 0.849 | 0.615 | 4.073 | 0.246 |

\* AUC: area under the curve; CI, confidence interval; LR+, positive likelihood ratio; LR-, negative likelihood ratio.

**Figure 5.**
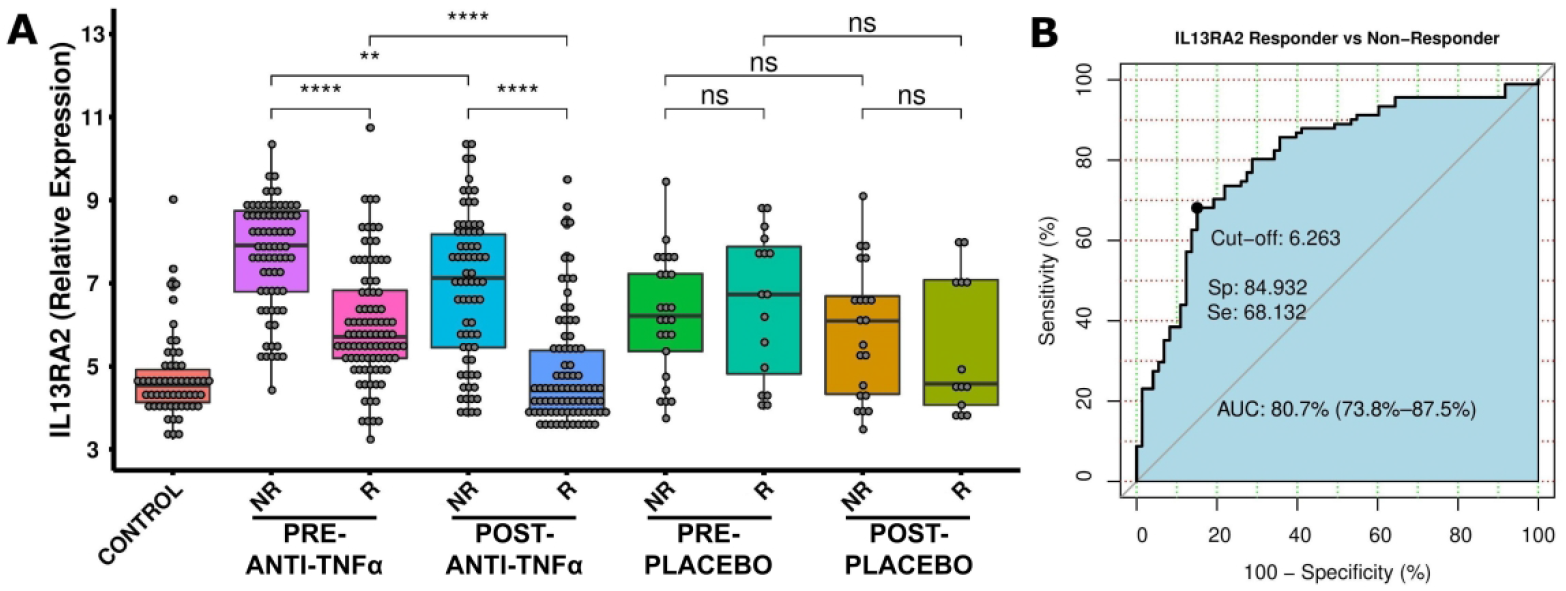
Interleukin 13 receptor subunit alpha 2 (IL13RA2) can be a diagnostic biomarker to predict TNFα treatment Response. (A) The expression of IL13RA2 is significantly higher in the pre-TNFα non-responders compared to the pre-TNFα responders. (B) The expression of IL13RA2 has an AUC of 80.7% (73.8% - 87.5%) with a sensitivity of 68.13% and specificity of 84.93%. NR: non-responder; R: responder; AUC: area under the curve, Sp: Specificity, Se: Sensitivity. Asterisks denote statistically significant differences (**p < 0.01, ***p < 0.001, ***p < 0.0001). Statistical significance was determined by 2-tailed Student’s t-test.

## Discussion

Treatment resistance of anti-TNFα is a critical issue in IBD patients. By integrating the existing raw data and increasing the statistical power, we revealed that immune microenvironment scores are higher in treatment resistance patients on baseline level (**Figure 1A–B**), indicating a higher inflammatory burden in anti-TNFα treatment non-responders. Further in-depth analysis uncovered neutrophils, endothelial and B cells contribute to the changes of the inflammatory burden (**Figures 3**). Next, a total of 64 up-regulated genes were identified (**Figure 4A–B**) and neutrophil chemotaxis (4 out of the top 12 enrichment terms) may contribute to anti-TNFα treatment resistance in IBD patients (**Figure 4D–E**). Utilising an animal study model, mean expression level (across samples) of genes matching the four chemotaxis GO terms are upregulated in LPS-induced neutrophils but no statistical changes in the adalimumab-treated group (**Figure 4**).

In a typical inflammatory response, immune cells such as macrophages, dendritic cells, natural killer cells, and T lymphocytes release TNFα proinflammatory cytokines, leading to the activation of endothelial cells and neutrophils.^32^ The activation of endothelial cells in colonic mucosa enhances vascular permeability and induces the recruitment of immune cells, leading to the activation of chemotaxis. The activation of neutrophils follows the tethering, rolling, crawling and transmigration process from the blood vessel into the inflamed colonic tissues.^33^ When neutrophils engulf invasive gut microbiome, they release granule proteins and chromatin to form neutrophil extracellular traps (NETs) and secrete anti-microbial peptides to mediate extracellular killing of microbial pathogens.^34^ However, hyperactive neutrophils trigger an unrestrained activity of the positive feedback amplification loops, leading to endothelial cells and the surrounding tissues damage, inducing resolution delay (IL6, TNF*α*, and IFN*γ*) and chemokines (IL8, CCL3, and CCL4), which further the recruitment of neutrophils, monocytes and macrophages to the inflamed sites.^35^ The use of anti-TNFα blockers significantly suppress the infiltration of neutrophils and B cells population in the inflamed mucosa, and suppresses proinflammatory mediators, such as calprotectin (S100A8/A9), IL8, IL6, and TNFα production,^36,37^ and matched with our finding only in responders (**Figure 2A–C and Figure 2G–I**). The unwanted immunogenicity, however, has a high level of B cells due to the presence of anti-drug antibodies (ADAs).^38^ The presence of ADAs neutralise, interfere and/or alter the binding efficacy, as well as pharmaco-dynamic/-kinetic properties of anti-TNFα monotonical antibodies.^39^

There are several S100 calcium-binding protein family genes that are highly expressed and previously studied in anti-TNFα treatment (**Figure 3A–B**). Calprotectin is a calcium-binding protein from the S100A8 and S100A9 monomers, representing up to 40% of neutrophil cytosolic proteins and constantly released from the inflamed region(s).^40^ *S100A12*, also known as calgranulin C, is released from neutrophils^40^ and participates in proinflammatory process via the activation of the NF-κB.^41^ A Small-scale study reported that faecal calprotectin test (commonly used to distinguish between irritable bowel syndrome (IBS) and IBD) and *S100A12* may predict relapse after one year of infliximab treatment^42^ while another faecal calprotectin study did not find the difference.^43^

*IL13RA2* is stand alone in the volcano plot with the highest fold change and the lowest p-value (**Figure 3B, Supplementary Data 4**), and has the best AUC (80.7%, 95% CI: 73.8% - 87.5%) outcome (**Table 2, Figure 5B, Supplementary Data 5**). Early studies uncovered *IL13RA2* is active in mucosal biopsies on the UC or CD anti-TNFα treatment non-responders compared to the responders.^44,45^ A small scale study demonstrated soluble IL13RA2 protein cannot be detected in serum, and tissue expression of IL13RA2 could predict anti-TNFα treatment in CD patients.^46^ IL13RA2 is a decoy receptor enable to bind IL13 cytokine, diminishes its JAK1/STAT6-mediated effector functions and activates activator protein 1 (AP-1) to induce the secretion of TGF-β.^47,48^ The IL13 pathway is also dependent on the production of TNFα. Several IL13 targeting drugs have been tested to inhibit hyperactive immune response on Th2-driven inflammatory diseases.^48^ However, insufficient protection was demonstrated by the phase IIa Anrukinzumab (an IL13 monoclonal antibody) clinical trial on UC patients.^49^ Thus, blocking the IL-13 pathway via IL13RA2 could be a new approach in treating IBD patients. IL13RA2 knock-out mice in DSS induced acute colitis model showed a better recovery rate compared to the wide-type mice, and negatively regulate epithelial/mucosal healing.^50^ By neutralising IL13RA2 in DSS induced IBD murine model using a monoclonal antibody, it presented a speedy recovery compared to the control group.^47^

The study here identified many strengths but should be considered in the context of shortcomings. Firstly, we only focused on data from large intestine and eliminated ileum data from the Arijs *et al*. study due to the low number of ileum samples that can be integrated,^27^ and also reduce the gene expression variation between two different organ sites for down-stream analysis. Secondly, our comparison does not include studies from vedolizumab and ustekinumab as it has limited datasets available online. Thirdly, the anti-TNFα response criteria and the determination time points are slightly different between studies. As we can only rely on the information provided by the authors, and thus our study has to accept the potential bias. Fourthly, anti-TNFα is broadly used in colitis-based diseases with a high percentage of treatment failure, and the diagnosis criteria of UC/CD/IBDU on each of the included studies may be slightly different with a certain percentage of misclassification.^10^ Therefore, our priority is to find the common patterns to minimise the treatment resistance rate in this study. Last but not least, the animal study in neutrophils are not from the colonic tissue sites and some of the chemotactic factor markers such as IL8/CXCL8 and CSF3 have a significant reduction after the adalimumab treatment.^26^ We believed that some single markers may not represent a whole picture of chemotaxis. Thus, in the early future, IBD subtypes analysis and more in-depth study in the relation to hyperactive chemotaxis are needed.

In conclusion, pre-anti-TNFα treatment non-responder patients presented a higher population of neutrophils, endothelial and B cells compared to the responders and the responders suppressed the activity of the immune cells. *IL13RA2* is a potential biomarker to predict anti-TNFα treatment response.

## Ethics approval

No ethical clearance required because this study retrieved and synthesised the data from already published studies, and researchers of each of the original studies obtained approval from their local ethics committee.

## Consent to participate

Not available

## Consent for publication

Not available

## Availability of data and materials

All the transcriptomics data were retrieved from publicly available datasets from the NCBI Gene Expression Omnibus (GEO) database. All the *in silico* cell sorting algorithms used in this study were based on the default settings recommended by the corresponding authors, either from web-based or retrieved from the corresponding authors’ GitHub page for academic research purposes.

## Authors’ contributions

Study conception and design: Tung On Yau; Development of methodology: Tung On Yau; Data acquisition and analysis: Tung On Yau, Guodong Du; Data interpretation: Tung On Yau, Jayakumar Vadakekolathu, Sergio Rutella. Writing - original draft preparation: Tung On Yau; Writing - review and editing: Jayakumar Vadakekolathu, Gemma Ann Foulds, Benjamin Dickins. Administrative, technical or material support: Benjamin Dickins, Christos Polytarchou, Sergio Rutella. Study supervision: Christos Polytarchou, Sergio Rutella.

## Acknowledgement

Not applicable

---

**Supplementary Figure 1.**
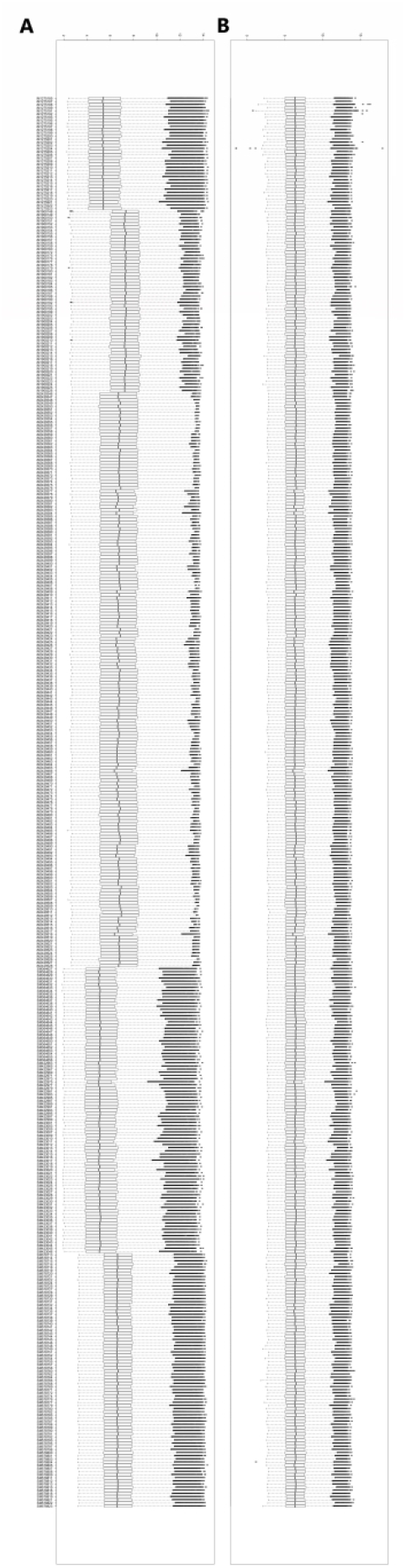
(A) Before and (B) after the batch effects correction of the included samples.

**Supplementary Figure 2.**
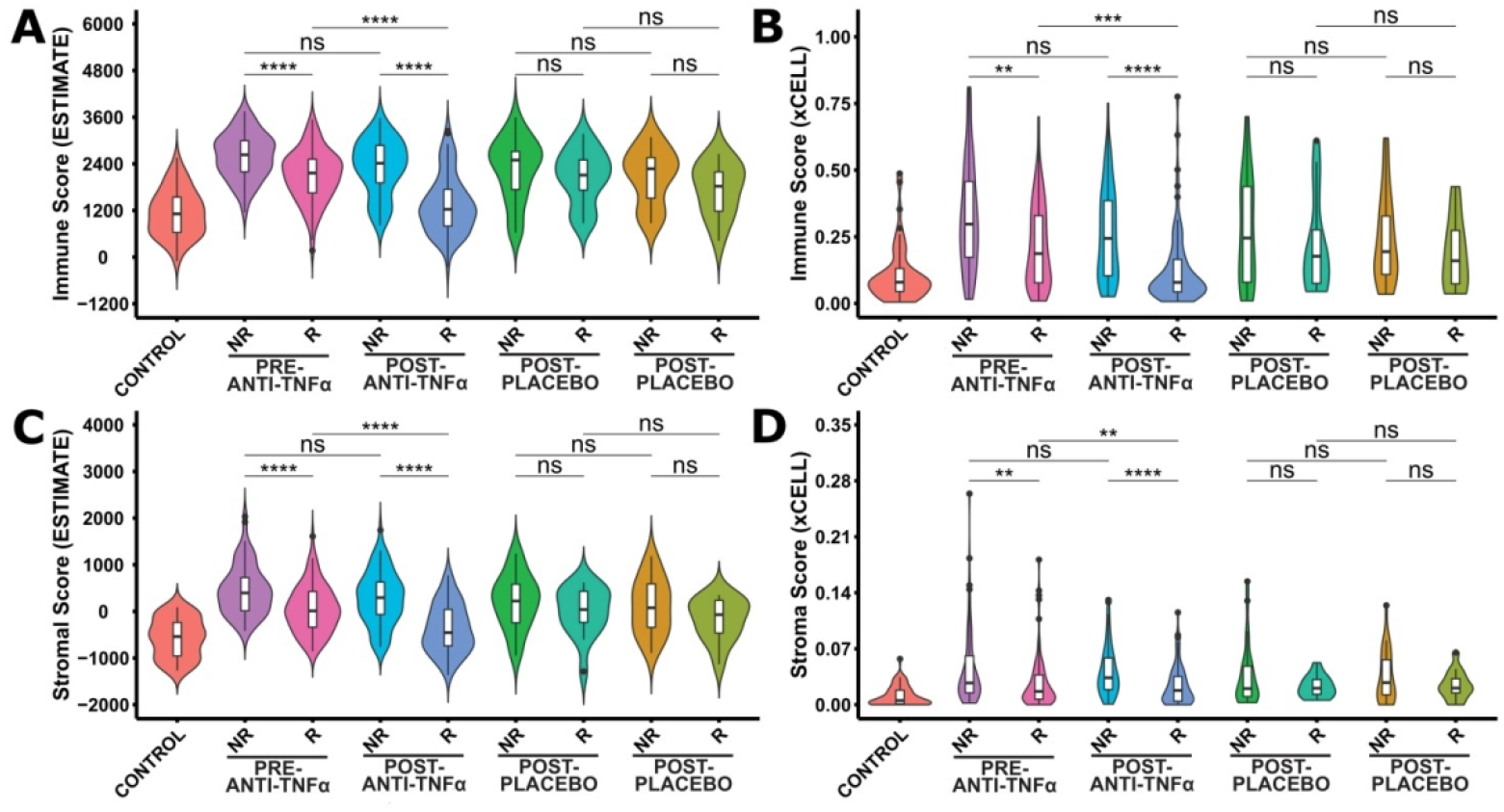
Immune-Stroma scores are significantly higher on baseline non-responders compared to the responders. Scores evaluated using both xCELL and ESTIMATE algorithms for (A, B) immune and (C, D) stroma/stromal scores are significantly higher in the pre-anti-TNFα treatment non-responders compared to responders. NR: non-responder; R: responder. The y-axes are the relative immune microenvironment scores from the corresponding algorithms. P-value determines by Mann-Whitney test with Benjamini and Hochberg adjustment. Asterisks denote statistically significant differences (*p < 0.05, **p < 0.01, ***p < 0.001, ****p < 0.0001).

**Supplementary Figure 3.**
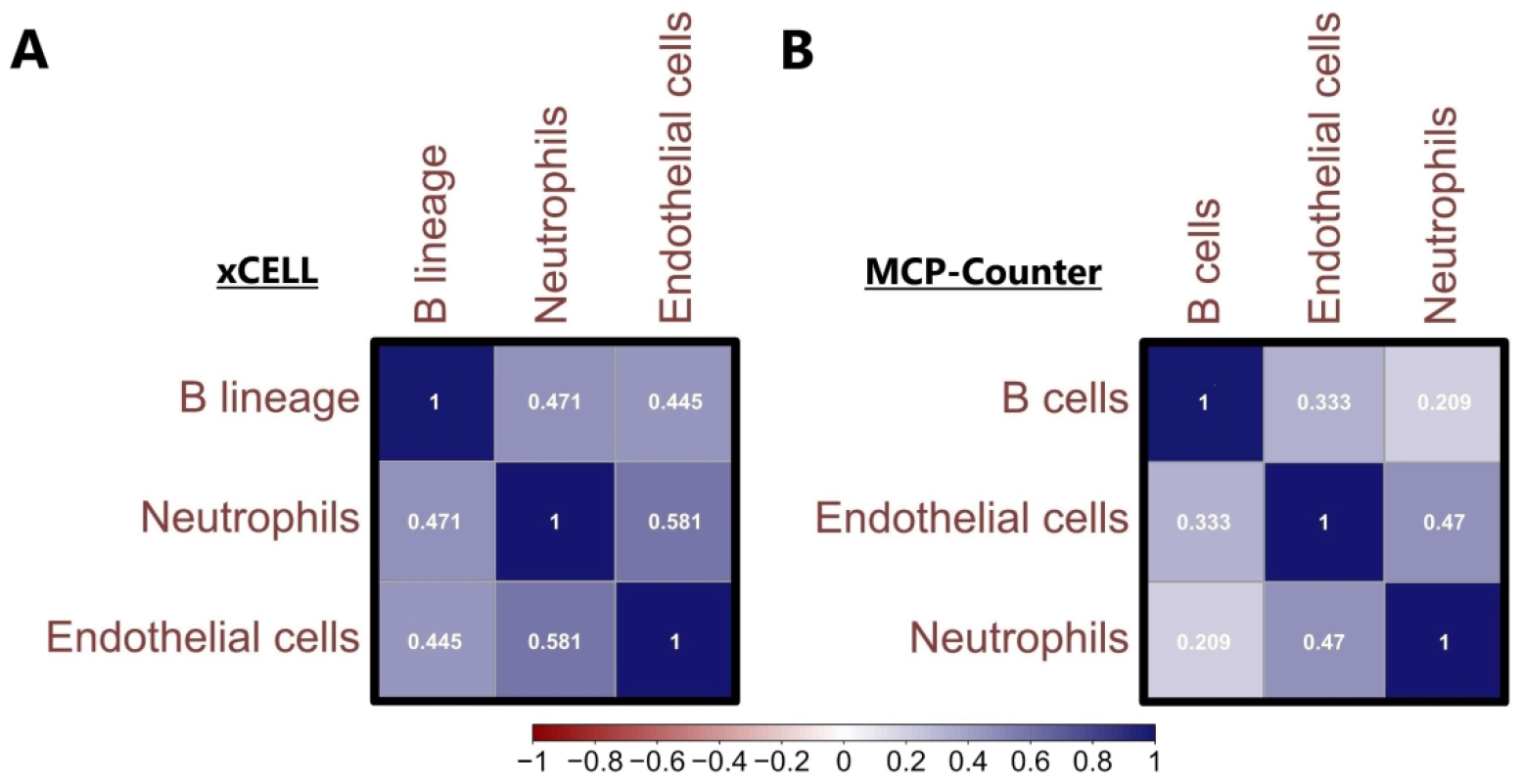
Pearson correlation matrix of Neutrophils, Endothelial cells and B cells from (A) xCELL and (B) MCP-Counter algorithms. Blue indicates positive correlation, and red indicate negative correlation. Darker colours are associated with stronger correlation coefficients.

**Supplementary Figure 4.**
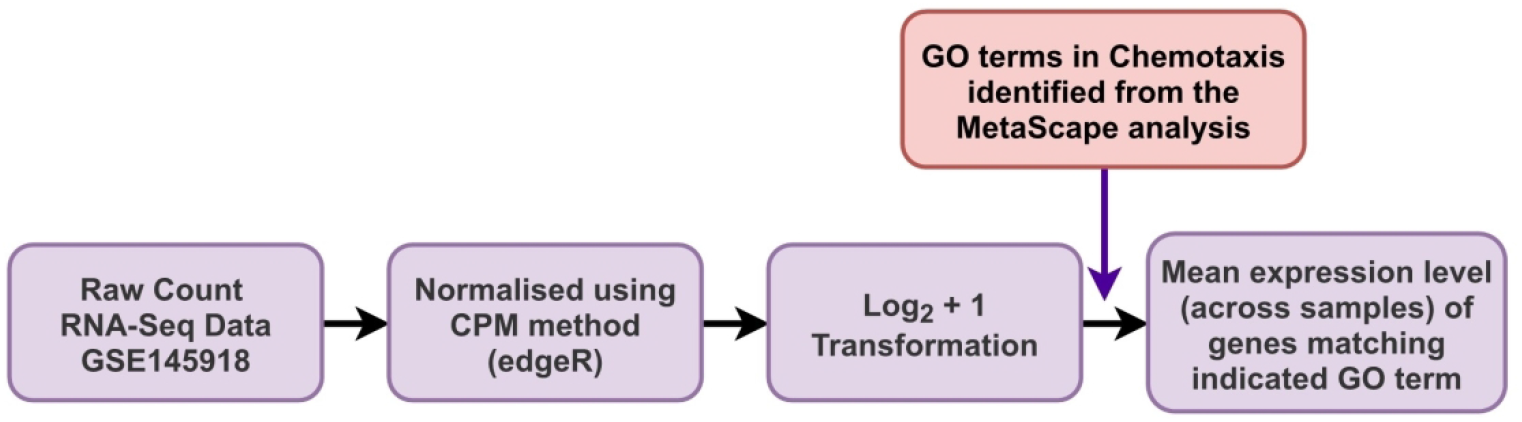
The RNA-sequencing data process workflow in LPS-induced inflammation in neutrophils.

